# Hypothalamic estrogen receptor alpha mediates key side effects of tamoxifen therapy in mice

**DOI:** 10.1101/2020.09.21.307124

**Authors:** Z Zhang, J. W. Park, I. S. Ahn, G. Diamante, N. Sivakumar, D. V. Arneson, X. Yang, J. E. van Veen, S. M. Correa

## Abstract

Adjuvant tamoxifen therapy for invasive breast cancer improves patient survival. Unfortunately, long-term treatment comes with side effects that impact health and quality of life, including hot flashes, changes in bone density, and fatigue. Partly due to a lack of proven animal models, the tissues and cell types that mediate these negative side effects are largely unknown. Here we show that mice undergoing a 28-day course of tamoxifen treatment experience dysregulation of core and skin temperature, changes in bone density, and decreased physical activity, recapitulating key aspects of the human physiological response. Single cell RNA sequencing reveals that tamoxifen treatment induces significant and widespread gene expression changes in different cell types of the hypothalamus, most strongly in neurons and ependymal cells. These expression changes are dependent on estrogen receptor alpha (ERα), as conditional knockout of ERα in the hypothalamus ablated or reversed tamoxifen-induced gene expression. Accordingly, ERα-deficient mice do not exhibit changes in thermal regulation, bone density, or movement in response to tamoxifen treatment. These findings provide mechanistic insight into the effects of tamoxifen on the hypothalamus and support a model in which hypothalamic ERα mediates several key side effects of tamoxifen therapy.

## Introduction

Tamoxifen is a selective estrogen receptor modulator that has been used for effective treatment of hormone responsive breast cancers for more than 40 years *(Jordan 2003)*. As an adjuvant, tamoxifen therapy can decrease the incidence of breast cancer recurrence by up to 40 percent *(Davies et al. 2011)*. This exceptionally effective treatment remains standard of care for people with hormone-responsive cancers, and reduction of recurrence persists for at least ten years of continuous tamoxifen treatment *(Davies et al. 2013, Chlebowski et al. 2014, Gierach et al. 2017)*. In contrast to these benefits, tamoxifen treatment has been associated with a variety of negative side effects including increased risk for hot flashes*(Love et al. 1991, Howell et al. 2005, Francis et al. 2015)*, endometrial cancer, venous thromboembolic events *(Fisher et al. 1998, Cuzick et al. 2007)*, bone loss *(Powles et al. 1996)*, and fatigue *(Haghighat et al. 2003)*. These responses markedly impact quality of life. Accordingly, ∼25% of eligible patients fail to start or complete this life-saving therapy due to side effects and safety concerns *(Friese et al. 2013, Berkowitz et al. 2020)*. The tissues and cells that mediate these negative side effects remain unclear. Unraveling the cells and mechanisms that mediate the positive effects of tamoxifen from those that mediate the negative side effects is necessary for understanding the multifaceted effects of tamoxifen therapy on physiology. Ultimately, this knowledge could lead to the design of new or adjuvant therapies that circumvent the side effects, improve patient quality of life, and perhaps enhance survival via increased patient compliance.

Within the brain, the hypothalamus is highly enriched for estrogen receptor expression and represents an excellent anatomical candidate for mediating many of the side effects of tamoxifen therapy. Estrogen receptor alpha (ERα) signaling regulates body temperature *(Bowe et al. 2006, Musatov et al. 2007, Mittelman-Smith et al. 2012, Martinez de Morentin et al. 2014)*, physical activity *(Musatov et al. 2007, Correa et al. 2015, van Veen et al. 2020)*, and bone density *(Farman et al. 2016, Zhang et al. 2016, Herber et al. 2019)* through distinct neuron populations in the hypothalamus. Indeed, the hypothalamus is a demonstrated target of tamoxifen, leading to changes in food intake and body weight *(Wade et al. 1993, Lopez et al. 2006, Lampert et al. 2013)* and changes in the hypothalamic-pituitary-ovarian *(Wilson et al. 2003, Aquino et al. 2016)* and hypothalamic-pituitary-adrenal *(Wilson et al. 2003)* axes. Tamoxifen has also been shown to affect gene expression in the hypothalamus; its administration blocks the estrogen dependent induction of the progesterone receptor (*Pgr*) in the ventromedial hypothalamus (VMH) and increases the expression of estrogen receptor beta (*Esr2*) in the paraventricular nucleus of the hypothalamus (PVH) *(Patisaul et al. 2003, Aquino et al. 2016, Sa et al. 2016)*.

We hypothesized that tamoxifen alters estrogen receptor signaling in the hypothalamus to mediate key negative side effects of tamoxifen therapy. To test this hypothesis, we modeled tamoxifen therapy in mice with a 28-day treatment course based on human dosage *(Slee et al. 1988)* and asked if mice experience physiological effects similar to humans. We measured physical activity, bone density, and the temperature of the body core, tail skin, and thermogenic brown adipose tissue (BAT). Profiling genome-wide expression changes of individual cells in the hypothalamus using Drop-seq, a droplet-based single-cell RNA sequencing technology, revealed transcriptional changes induced by tamoxifen in multiple cell types. Finally, we show that hypothalamic ERα signaling is necessary for both the gene expression changes in the hypothalamus and the effects on thermoregulation, bone density, and physical activity. Together, these findings suggest that tamoxifen therapy modulates hypothalamic ERα signaling to alter fundamental aspects of physiology and health. Dissecting central versus peripheral effects and mechanisms of tamoxifen therapy is the first step toward identifying strategies to mitigate the adverse side effects of this life-saving treatment.

## Results

### Tamoxifen treatment alters thermoregulation

To ask if mice and humans experience similar physiological effects while under tamoxifen treatment, we administered tamoxifen (0.1mg/kg) or vehicle subcutaneously, daily, for 4 weeks (Figure 1A). In humans, hot flashes are characterized by frequent and sudden increases of heat dissipation from the face and other parts of the skin, often accompanied with perspiration and a decrease in core body temperature *(Stearns et al. 2002)*. Similarly, tamoxifen treated mice showed significantly lower core body temperature compared to controls, as indicated by 24-hour averages from the last week of treatment. This difference is detected during the light phase when the animals are generally inactive but not in the dark phase when physical activity can also influence core temperature (Figure 1B and C). In mice, heat dissipation occurs effectively via vasodilation in the tail *(Gordon 1993)*. As tail skin temperature is highly dependent on core and ambient temperature, heat loss is often expressed as the heat loss index (HLI): HLI = (Tskin − Tambient)/(Tcore − Tambient) *(Romanovsky et al. 2002)*. We observed a higher HLI in mice treated with tamoxifen compared to controls (Figure 1D). Again, this effect was detected in the light phase but not in the dark phase (Figure 1E). Tail skin temperature also was significantly higher in tamoxifen treated mice compared to controls during light phase (Figure 1 – Supplement 1A and B). In addition, mice treated with tamoxifen exhibited lower temperature above the intrascapular region directly apposed to BAT depots (Figure 1F and G), suggesting reduced heat production from BAT following tamoxifen treatment. Accordingly, postmortem qPCR analysis of BAT revealed lower expression of genes associated with thermogenesis, uncoupling protein 1 (*Ucp1*) and adrenergic receptor beta 3 (*Adrb3*) (Figure 1 – Supplement 1C), suggesting suppressed BAT thermogenesis and sympathetic tone following tamoxifen treatment *(Cannon et al. 2004)*. Together, these results indicate that tamoxifen treatment shifts mouse temperature balance toward increased heat dissipation and decreased heat production, consistent with the observations in humans experiencing hot flashes.

**Figure 1.**
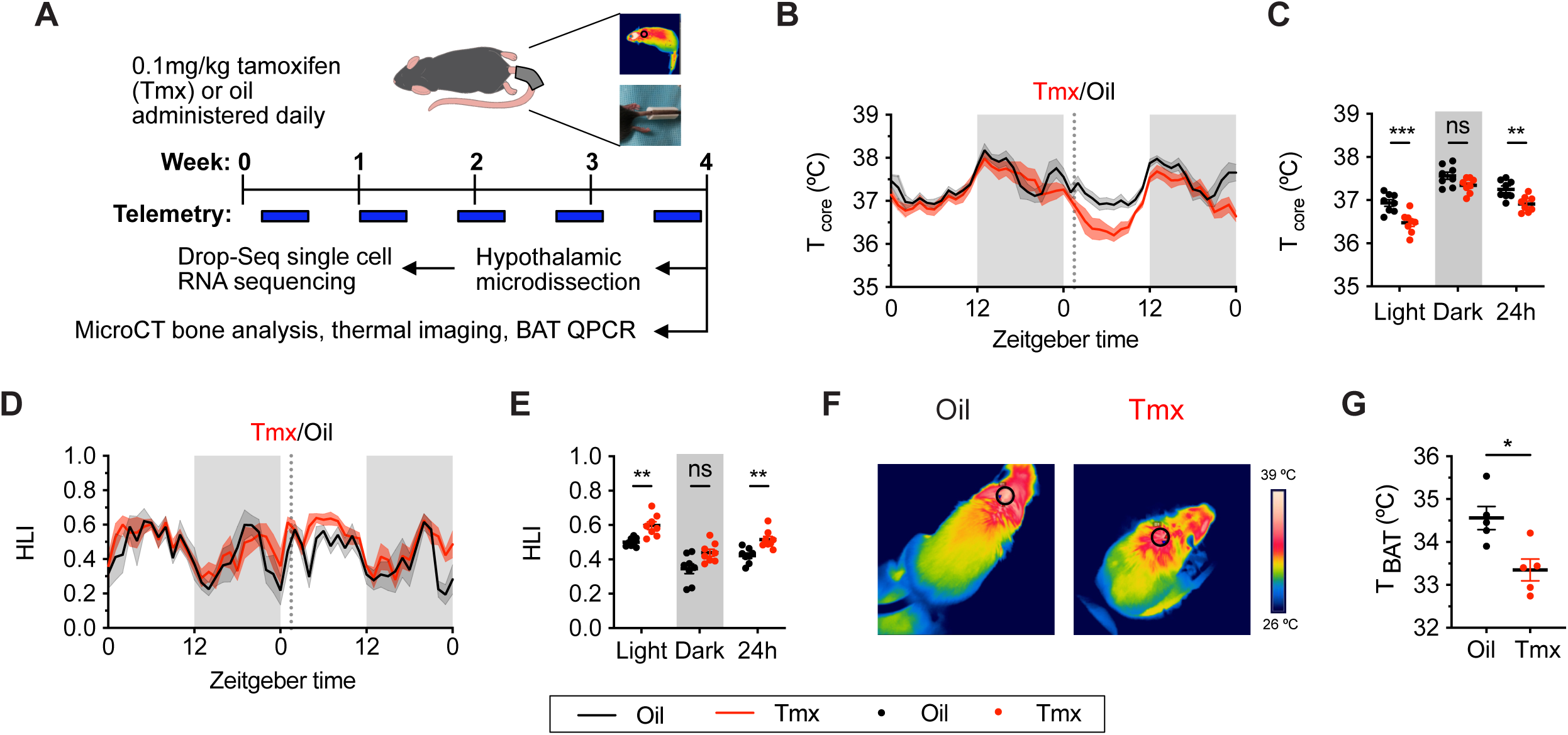
Tamoxifen treatment decreases core body temperature and increases heat dissipation. (**A**) Strategy to measure the physiological and gene expression effects of long term tamoxifen (Tmx) treatment. Mice were treated by daily subcutaneous injection of 0.1mg/kg tamoxifen or corn oil for 4 weeks. Core and tail temperature were measured every 5 minutes using telemetry probes. Snap shot of thermographic images for BAT temperature was obtained in last week of treatment. At the conclusion of 4 week treatment, experimental mice were sacrificed for bone analysis or single cell RNA sequencing. (**B**) Hourly average of core body temperature over 3 days before and 3 weeks after injections (dotted line). (**C**) Average core body temperature from mice shown in panel **B** highlighting per animal averages in light (7:00 to 19:00), dark (19:00 to 7:00), and total 24 h periods (n=8, treatment **) (**D**) Heat loss index (HLI) calculated from continuous measurements of core and tail temperature. (**E**) Average HLI from mice shown in panel **D** (n=7, treatment **). (**F**) Thermographic images of interscapular skin above brown adipose tissues depots in mice injected with either oil or tamoxifen. (G) Quantification of temperature of skin above interscapular BAT depots (n=5). In **B** and **D**, line shading width represents SEM. *, P<0.05; **, P<0.01; ***, P<0.001 for Sidak’s multiple comparison tests (**C** and **E**) following a significant effect of treatment in a two-way RM ANOVA or two tailed t-tests (**G**).

### Tamoxifen treatment increases bone density and decreases movement

Tamoxifen has been shown to affect bone density in rodent and human studies *(Powles et al. 1996, Perry et al. 2005)*. Here, micro CT scans revealed that tamoxifen treatment is associated with greater bone volume ratios in both metaphysis and diaphysis of the femur (Figure 2A-C). Tamoxifen treatment was also associated with greater number and thickness, and less separation of trabecular bones in metaphysis (Figure 2B). Bone mineral density and bone area in cortical bones were not significantly affected by tamoxifen treatment (Figure 2C); however, tamoxifen treatment was associated with significantly greater cortical thickness (Figure 2A and C). Together, these data indicate that chronic tamoxifen treatment increases bone mass, consistent with previous studies showing increased bone formation and bone mass in mice following tamoxifen administration *(Perry et al. 2005, Starnes et al. 2007)*. In addition, tamoxifen resulted in a moderate decrease in 24 h physical activity (Figure 2D and E). Tamoxifen treated mice did not show significant changes in body weight compared to control mice, over 28 days of treatment (Figure 2F and G).

**Figure 2.**
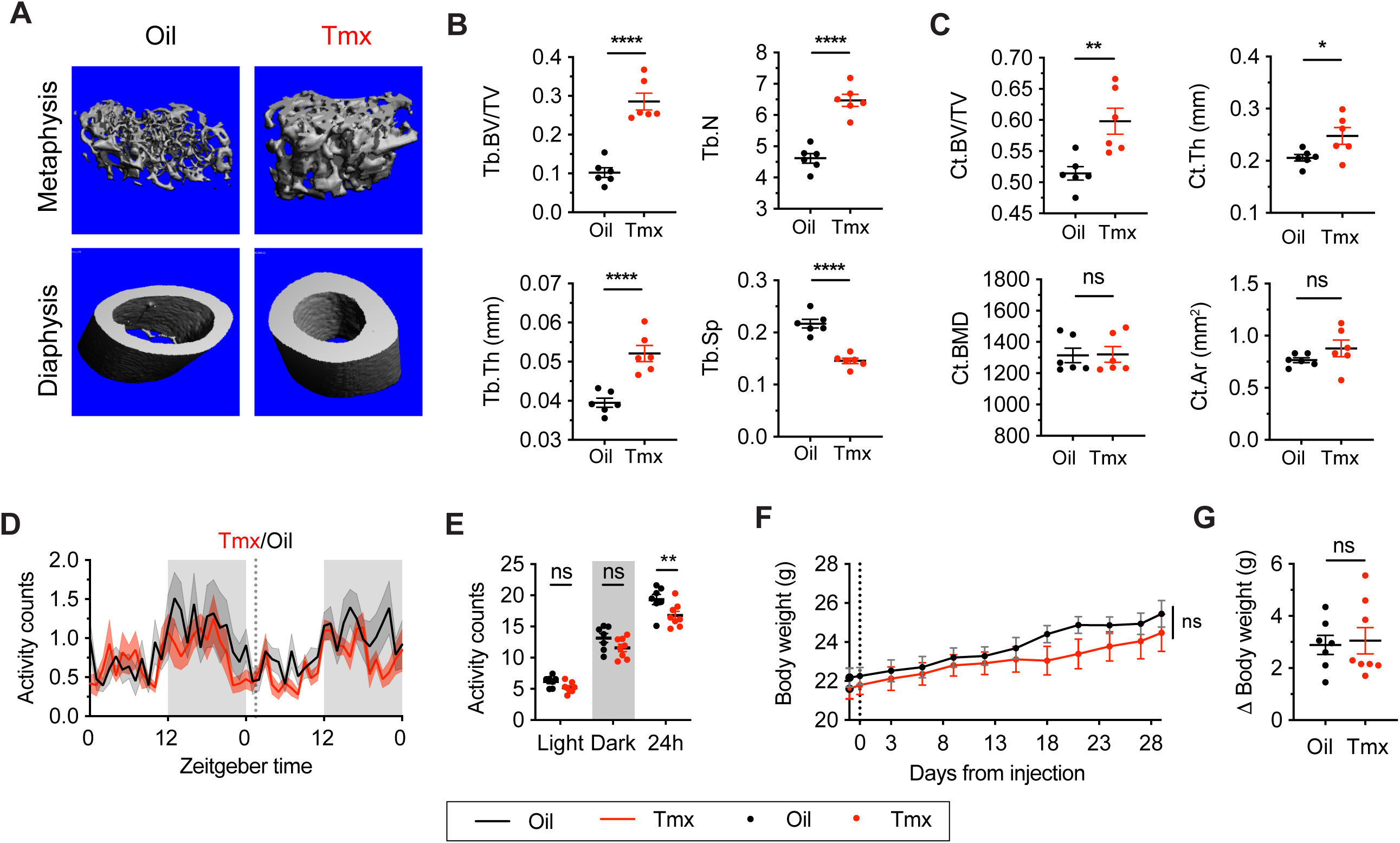
Tamoxifen treatment increases bone density and decreases movement. (**A**) Representive microCT images showing bone density of the distal metaphysis and midshaft diaphysis of femurs from tamoxifen or oil treated female mice. Tamoxifen or vehicle was injected subcutaneously at 0.1mg/kg for 28 days. (**B**) Trabecular bone volume fraction (BV/TV), trabecular numbers (Tb.N), trabecular thickness (Tb.Th) and trabecular separation (Tb.Sp) in distal metaphysis of femurs (n=6). (**C**) Bone volume fraction (Ct.BV/TV), cortical thickness (Ct.Th), bone mineral density (Ct.BMD) and cortical area (Ct.Ar) in diaphysis of femurs (n=6). (**D**) Hourly average of physical activity over 3 days before and 3 weeks after tamoxifen or oil injections (dotted line) measured every 5 min using intraperitoneous telemetry probe. Line shading width represents SEM. (**E**) Average total physical activity from mice shown in panel **D** (n=8, treatment *). (**F**) Change of body weight over 28 days during tamoxifen or oil injection (n=7-8, treatment P=0.5468). Error bars represent SEM. (**G**) total change in body weight over the course of the 28 day experiment (n=8, P=0.8010). ns, non-significant; **, P<0.01; ***, P<0.001; ****, P<0.0001 for Sidak’s multiple comparison tests (**E**) following a significant effect of treatment in a two-way RM ANOVA or two tailed t-tests (**B, C** and **G**).

### Tamoxifen administration affects gene expression in all hypothalamic cell types

To ask how tamoxifen affects gene expression in the different cell types of the hypothalamus, we collected hypothalami (Figure 3 – Supplement 1A) after 28 daily injections of tamoxifen or vehicle and analyzed individual cells by drop-Seq *(Macosko et al. 2015)*. A total of 29,807 cells (8,220 from n = 3 vehicle treated mice, 21,587 from n = 5 tamoxifen treated mice) clustered into 9 distinct cell types, annotated based on high expression of previously characterized cell type markers *(Moffitt et al. 2018, Saunders et al. 2018)*: astrocytes, oligodendrocytes, neurons, endothelial cells, microglia, polydendrocytes, ependymal cells, mural cells, and fibroblasts (UMAP: Figure 3A, tSNE: Figure 3 – Supplement 1B, cluster defining marker expression: Figure 3 – Supplement 1D). Because tamoxifen modulates estrogen receptor signaling *(de Médina et al. 2004, Jordan 2007)*, we examined cell-type specific expression of three estrogen receptor transcripts: the nuclear receptors *Esr1* and *Esr2*, as well as the G-protein-coupled estrogen receptor 1 (*Gper1*, formerly *Gpr30*) *(Revankar et al. 2005)*. While neither *Esr1, Esr2*, or the *Esr1* target gene *Pgr* showed strong enrichment in any particular cell type (Figure 3B), *Gper1* transcripts were predominantly found in Mural cells (LogFC: 1.39 compared to all other cell types, adj. P < 1e-128). These results are in accordance with the reported *Esr1* expression pattern in the hypothalamus *(Shughrue et al. 1997)* and *Gper1* in vascular smooth muscle of the rodent brain *(Isensee et al. 2009)*. The graphical clustering methods, UMAP and tSNE did not demonstrate clear separation between tamoxifen and vehicle treated cells (Figure 3C, Figure 3 – Supplement 1C), indicating that the effect of tamoxifen at clinically relevant doses may be modest with respect to global transcriptional signatures.

**Figure 3.**
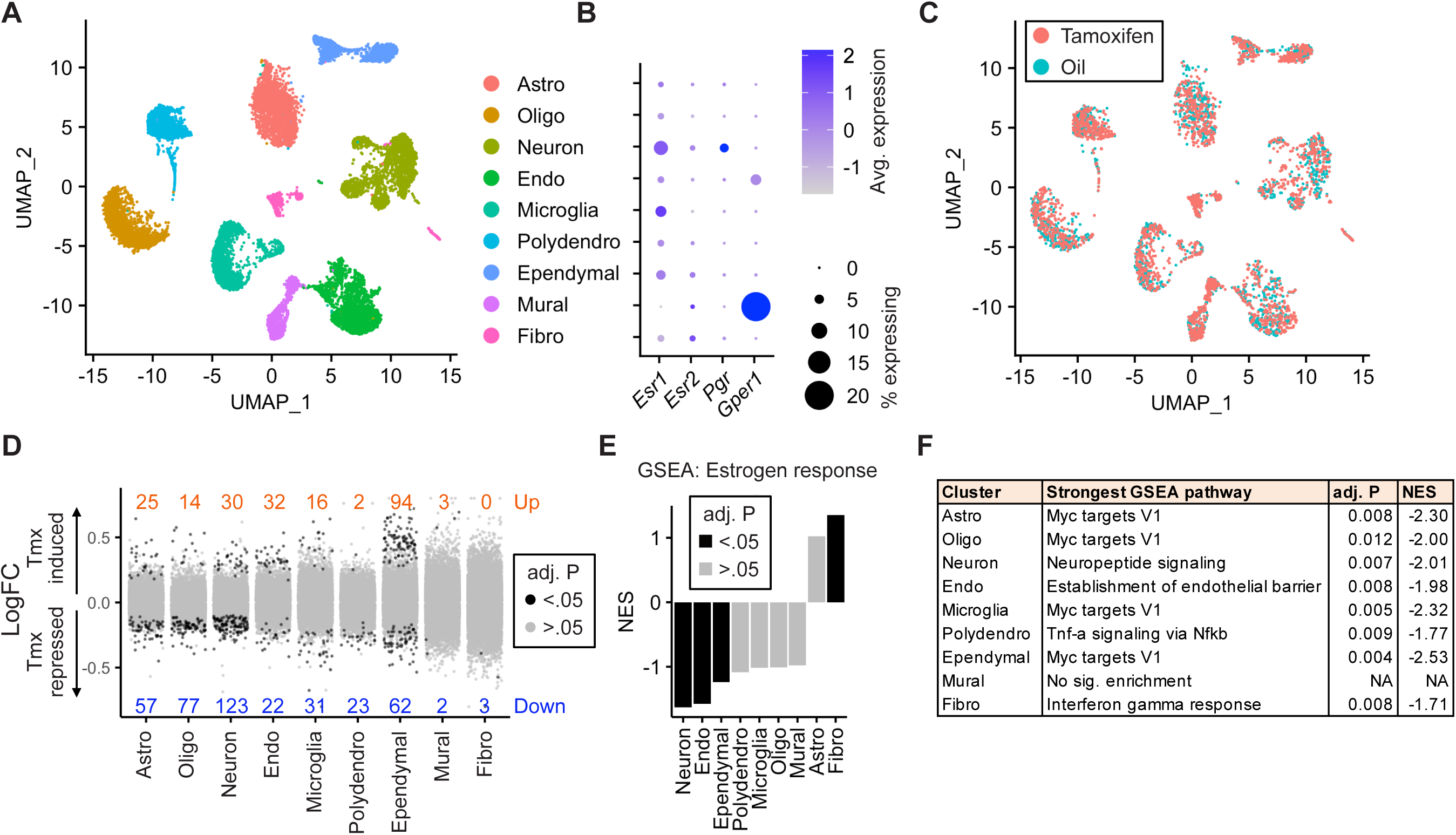
Tamoxifen treatment induces gene expression changes in various cell types of the hypothalamus. (**A**) UMAP showing clustering of the major cell types of the hypothalamus based on single cell transcriptomics. (**B**) Dot plot showing expression of estrogen and progesterone receptors in hypothalamic cell types. (**C**) UMAP of hypothalamic cells, comparing cells derived from female mice receiving daily tamoxifen injections (pink) or daily oil injections (blue). (**D**) Collapsed volcano plots showing differential gene expression, represented as log base 2 of fold change (LogFC), induced by daily tamoxifen treatment in various cell types of the hypothalamus. Up/Down numbers refer to total number of significantly (Bonferroni adj. P < .05) up and down regulated genes. (**E**) GSEA to find tamoxifen induced signatures of estrogen response in hypothalamic cell types. (**F**) Most strongly enriched or depleted pathways in GSEA comparing control to tamoxifen treated cells. All analyses done from cells harvested from female mice injected daily with oil (n=3) or tamoxifen (n=5) over 28 days.

Amongst all cell types, neurons and ependymal cells were most sensitive to tamoxifen administration, as indicated by the number of significantly induced and repressed genes (Figure 3D). To account for the effect of cluster size on statistical power, we also calculated the number of significant differentially expressed genes per 1k cells in each cluster. Again, neurons and ependymal cells have the highest numbers of differentially expressed genes when expressed as total or as a fraction of the number of cells sampled (Figure 3 – Supplement 1E), suggesting the strongest tamoxifen responsiveness. Although mural cells show significant enrichment of *Gper1*, tamoxifen administration was associated with relatively few significant gene expression changes in these cells (Figure 3D), indicating that *Gper1* does not likely mediate tamoxifen induced gene expression in the hypothalamus. It is possible, however, that tamoxifen affects cellular state in a non-transcriptional manner, via *Gper1*. Gene set enrichment analysis (GSEA) using a gene set for estrogen response (GO:0043627) further identifies neurons and ependymal cells as markedly tamoxifen responsive: tamoxifen administration induced a significant downregulation of genes involved in estrogen response in neurons, endothelial cells, and ependymal cells (Figure 3E, Table S1). Individual gene expression changes which may be of interest can be queried at https://correalab.shinyapps.io/tamoxifenshiny/.

GSEA with a collection of 40 hallmark pathways and brain-focused gene ontology gene sets (Table S2) revealed that all cell types except for mural cells showed significant enrichment or depletion of genes in at least one pathway (Figure 3F). Significantly enriched pathways (Table S3), show both overlapping and distinct effects by cell type. For example, tamoxifen treatment decreased the expression of genes annotated as targets of the proto-oncogene, *Myc*, in astrocytes, oligodendrocytes, neurons, endothelial cells, microglia, polydendrocytes, and ependymal cells. Other cell-type specific pathways were also enriched or depleted, including: establishment of the endothelial barrier in endothelial cells, neuropeptide signaling in neurons, E2F targets in astrocytes, and fatty acid metabolism in ependymal cells. Together these findings suggest that tamoxifen treatment has widespread effects on the hypothalamus, altering transcriptional programs and cell function in both general and cell-type specific ways.

### Tamoxifen treatment causes gene expression changes in neuronal sub-types of the hypothalamus

Given the transcriptional effects of tamoxifen treatment observed in neurons and the heterogeneity of estrogen-sensitive neurons within the hypothalamus, we next examined the effect of tamoxifen therapy within individual neuronal clusters. Neuronal sub-clustering resulted in 25 apparent hypothalamic neuronal sub-types (Figure 4A). The majority of clusters were marked by expression of genes previously demonstrated to delineate hypothalamic neuronal types (Figure 4B) *(Chen et al. 2017, Romanov et al. 2017, Kim et al. 2019, van Veen et al. 2020)*. Markers of neuronal subtypes include neuropeptides, transcription factors, and cellular signaling messengers. *Esr1* expression was observed in various neuronal subclusters (Figure 4B), but was significantly enriched in neurons expressing the neuropeptide precursor, *Tac2* (log2FC = 1.08, adj. P = 5.7e-98 compared to all other neuronal subclusters). GSEA demonstrates that overall, tamoxifen administration is associated with a significant downregulation of genes involved in neuropeptide signaling, a critical aspect of neuronal function, as well as the two principal components fueling the sizeable energy demands of neuronal metabolism: oxidative phosphorylation and glycolysis *(Yellen 2018)* (Figure 4C). Notably, it has previously been shown that estrogens can directly regulate these metabolic pathways in the brain *(Brinton 2008, Brinton 2009)*. When divided into sub-types, few statistically significant gene expression changes caused by tamoxifen were observed, likely due to loss of statistical power resulting from relatively few cells in each individual cluster (Figure 4 – Supplement 1A). To gain a preliminary view at the effect of tamoxifen on individual neuronal sub-types, we performed GSEA on all clusters; 14 of 25 neuronal types showed enrichment of at least one pathway (Table S4). Neurons marked by expression of *Tac2* showed significant enrichment or depletion of the most pathways (Figure 4 – Supplement 1B). Interestingly, examining individual pathways shows differential responses by neuronal subtype. For example, genes involved in oxidative phosphorylation appear downregulated by tamoxifen in neurons expressing *Foxb1*, but the same gene set is upregulated in neurons expressing *Tac2* (Fig 4 – Supplement 1B). These cell-type specific effects highlight the additional insights that can be revealed by scRNA-seq compared to overall averages determined by bulk tissue sequencing. Additionally, these results are compatible with the previous findings that tamoxifen can have different effects in different regions of the brain *(Wilson et al. 2003, Chen et al. 2014, Gibbs et al. 2014, Aquino et al. 2016, Sa et al. 2016)* and consistent with cell-type-specific effects of tamoxifen within hypothalamic neurons.

**Figure 4.**
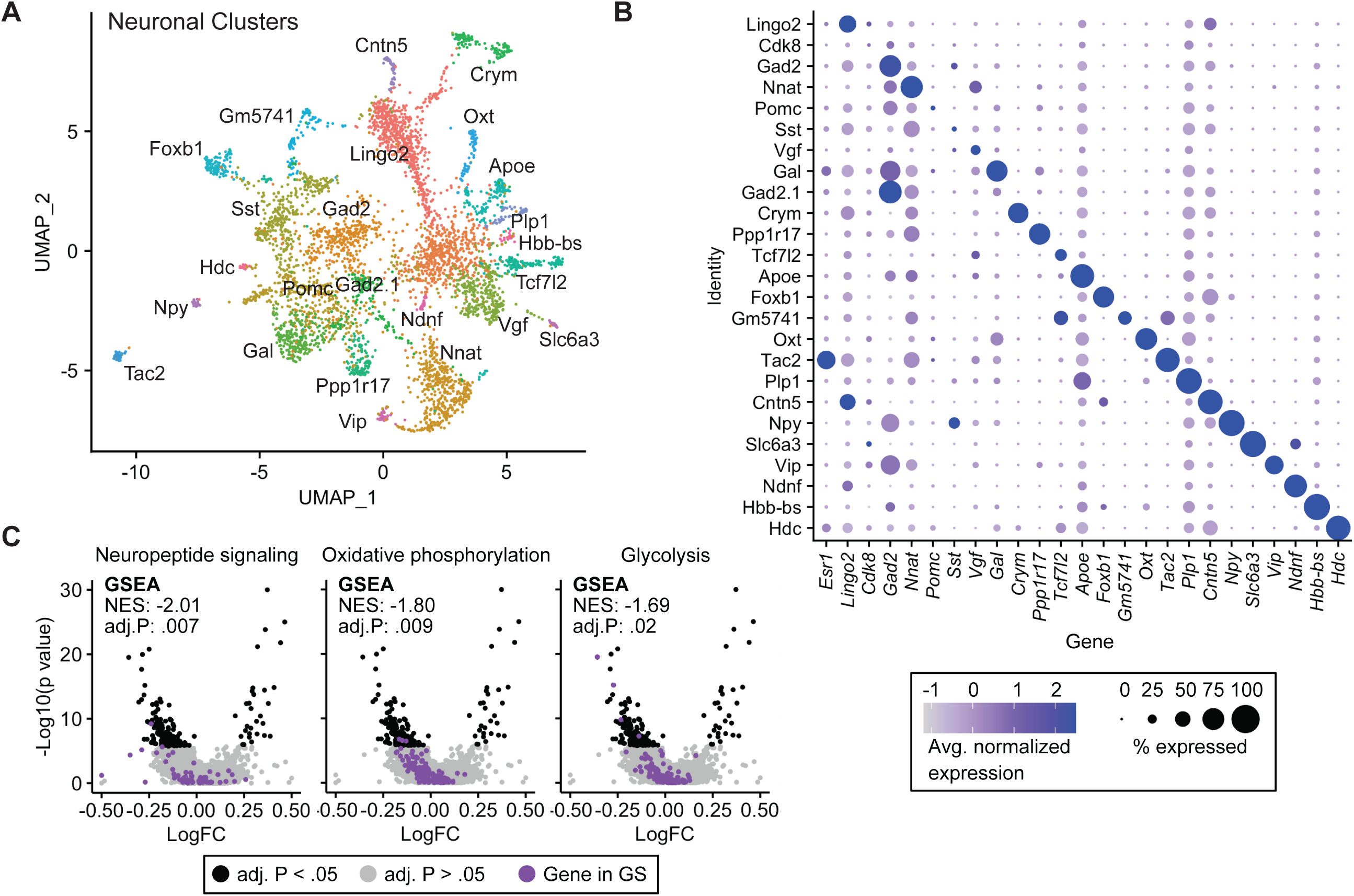
Tamoxifen induced gene expression changes in neurons of the hypothalamus. (**A**) UMAP showing clustering of the neuronal sub-types of the hypothalamus based on single cell transcriptomics, overlayed with identity named for top expressed cluster defining marker gene. (**B**) Dot plot showing expression of neuronal cluster defining markers and *Esr1*. (**C**) Volcano plots of tamoxifen induced or repressed DEGs overlayed with gene sets (GS) involved in neuropeptide signaling, oxidative phosphorylation, or Glycolysis. Analyses done from female mice injected daily with oil (n=3) or tamoxifen (n=5) over 28 days.

### *Esr1* knockout ablates hypothalamic gene expression responses to tamoxifen

As *Esr1* is a known target of tamoxifen that is expressed in the hypothalamus (Figure 3B), we sought to determine the impact of hypothalamic *Esr1* knockout on hypothalamic gene expression responses to tamoxifen administration. The transcription factor, *Nkx2-1*, is highly enriched in the medial basal hypothalamus and *Esr1*^*F/F*^; *Nkx2-1*^*Cre*^ (*Esr1*^cKO^) mice display a selective loss of ERα immunoreactivity in the hypothalamus *(Herber et al. 2019)*. Single-cell expression analysis was unable to detect a significant decrease of *Esr1* expression in any cell type of *Esr1*^cKO^ mice, but clearly demonstrated that the NK2 homeobox transcription factor *Nkx2-1* was enriched in neurons and in ependymal cells (Figure 5 – Supplement 2A). We also confirmed that *Esr1*^*F/F*^; *Nkx2-1*^*Cre*^ (*Esr1*^cKO^) mice displayed a loss of ERα immunoreactivity in the hypothalamus, including the medial preoptic area (MPA), ventrolateral subdivision of the ventromedial nucleus of the hypothalamus (VMHvl), and arcuate nucleus of the hypothalamus (ARC) (Figure 5A). These regions show highly enriched ERα expression *(Merchenthaler et al. 2004)* and mediate the primary hypothalamic effects of estrogens on body temperature *(Xu et al. 2011, Mauvais-Jarvis et al. 2013, Martinez de Morentin et al. 2014)*, physical activity *(Correa et al. 2015, van Veen et al. 2020)* and bone regulation *(Farman et al. 2016, Herber et al. 2019)*.

**Figure 5.**
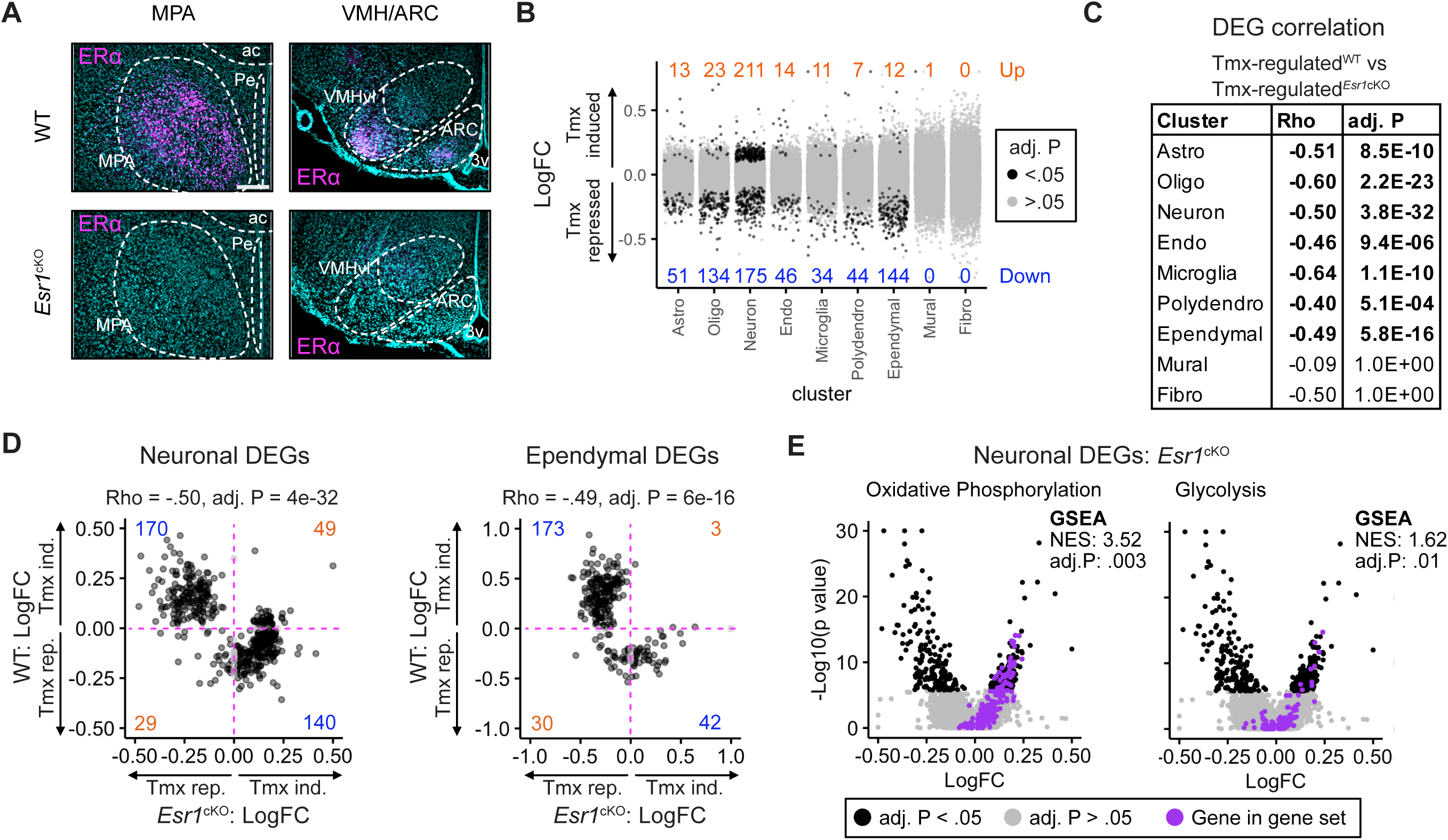
*Esr1* knockout reverses hypothalamic responses to tamoxifen. (**A**) Immunoreactive staining of ERα in the MPA, VMH, and the ARC of ERα knockout and wild-type female mice. Scale bar: 200 um. (**B**) Differential gene expression induced by daily tamoxifen treatment in cell types of the *Esr1*^*f/f*^;*Nkx2-1*^*Cre*^ (*Esr1*^cKO^) hypothalamus. Up/ Down numbers refer to total number of significantly (Bonferroni adj. P < .05) up and down regulated genes. (**C**) Table of correlations showing how tamoxifen induced gene expression changes in WT cells correlate with tamoxifen induced gene expression changes in *Esr1*^cKO^ cells. (**D**) Gene by gene comparison of how tamoxifen treatment affects expression in wild-type and *Esr1*^cKO^ neurons and ependymal cells. Blue numbers indicate ERα dependent DEGs, orange numbers indicate ERα independent DEGs (**E**) Volcano plots of all tamoxifen induced or repressed DEGs (black) overlayed with gene sets (purple) involved in oxidative phosphorylation or glycolysis. Data from n = 4 oil treated *Esr1*^cKO^ and n = 4 tamoxifen treated *Esr1*^cKO^, n = 3 oil treated WT and n = 5 tamoxifen treated WT mice.

To test the effect of tamoxifen administration on gene expression in mice lacking hypothalamic ERα, *Esr1*^cKO^ mice received the same daily tamoxifen or vehicle injection regimen concurrently with wild-type mice (Figure 1A). UMAP Clustering of all 4 treatment conditions: vehicle or tamoxifen in wild-type or *Esr1*^cKO^ animals did not reveal clear separation in global transcriptional signature between any treatment group when considering all cell types (Figure 5 – supplement 1A) or re-clustered neuronal sub-types (Figure 5 – Supplement 1B). *Esr1*^cKO^ animals showed proportionally fewer astrocytes and more oligodendrocytes than wild-type animals (Figure 5 – Supplement 1C), though all other cell types were recovered at similar proportions. Tamoxifen treatment was not associated with any significant differences in recovered cell type proportions (Figure 5 – Supplement 1D).

Interestingly, tamoxifen treatment led to significant transcriptional changes in the hypothalamus of *Esr1*^cKO^ mice (Figure 5B). Similar to wild-type, *Esr1*^cKO^ neurons and ependymal cells were the cell-types most sensitive to tamoxifen administration compared to all other cell types, as indicated by the number of significantly induced and repressed genes (Figure 5B). To look closer at the specific effects of tamoxifen on gene regulation, we examined only those gene expression changes that were significant in either wild-type or *Esr1*^cKO^ cells. When comparing how tamoxifen affected gene expression in wild-type mice versus how tamoxifen affected the same genes in *Esr1*^cKO^ mice, there was a strong negative Spearman correlation observed in all cell types, except the poorly powered mural cells and fibroblasts (Figure 5C). This negative correlation implies that the majority of significant tamoxifen-induced gene expression changes that were observed in hypothalamic cell types of wild-type female mice were ablated or regulated in the opposite direction in *Esr1*^cKO^ animals.

In wild-type neurons, the majority of tamoxifen-regulated genes were repressed by tamoxifen (Figure 3D). In *Esr1*^cKO^ neurons, most of these genes are no longer repressed by tamoxifen (Figure 5D, Key: Figure 5 – Supplement 2B). Together these data suggest that tamoxifen has generally suppressive effects on neuronal transcription, and that these effects are largely dependent on *Esr1*. In wild-type ependymal cells, the majority of regulated genes were induced by tamoxifen (Figure 3D). Again, this induction is largely dependent on the presence of ERα, as most genes induced by tamoxifen in wild-type ependymocytes were not induced by tamoxifen in *Esr1*^cKO^ ependymocytes. Despite displaying far fewer differentially expressed genes in either genotype, the other cell types showed similar trends (Figure 5 – Supplement 2C).

To determine if the overall opposite effects of tamoxifen in *Esr1*^cKO^ animals was associated with similar changes in functional pathways, we performed GSEA with all cell type clusters as before. Indeed, many pathways were altered in opposite directions in *Esr1*^cKO^ cells compared to wild-type cells following tamoxifen treatment (Complete results: Tables S2, S5). This pattern included *Myc* targets in endothelial cells, ependymal cells, neurons, oligodendrocytes, and polydendrocytes. Importantly, we also found a significant change of metabolic gene expression in neurons, as both oxidative phosphorylation and glycolysis were upregulated by tamoxifen in *Esr1*^cKO^ neurons (Figure 5E) but downregulated in wild-type neurons (Figure 4C). This trend of opposing effects of tamoxifen was not universal, as *Tnf-a* signaling was significantly downregulated by tamoxifen in both *Esr1*^cKO^ and wild-type endothelial cells. Finally, many pathways, including *mTorc1* signaling in astrocytes, establishment of the endothelial barrier in endothelial cells, and neuropeptide signaling in neurons, were significantly affected by tamoxifen in wild-type cells, but not affected in their *Esr1*^cKO^ counterparts.

GSEA for estrogen responsive genes in all cell types showed that tamoxifen downregulated estrogen targets in multiple cell types of the wild-type hypothalamus but did not significantly affect the same genes in cells of the *Esr1*^cKO^ hypothalamus, even when using a promiscuous cutoff or significance (Figure 5 – Supplement 3A, Table S1, S6). Similarly, GSEA in individual neuronal subclusters showed that tamoxifen broadly downregulated estrogen targets in wild-type neuronal subtypes, and while we still observed some downregulation of estrogen responsive genes in neuronal subtypes, there was never a significant upregulation of estrogen responsive genes by tamoxifen, even when using a permissive cutoff for significance (Figure 5 – Supplement 3B, Table S7, S8). Together these data indicate that hypothalamic expression of ERα is required for a large proportion of the gene expression changes normally induced by tamoxifen, including many functional pathways and those controlling neuronal activity and estrogen targets in many cell types and neuronal subtypes. Individual gene expression changes which may be of interest can be queried at https://correalab.shinyapps.io/tamoxifenshiny/.

### The homeostatic effects of tamoxifen are dependent on hypothalamic ERα

Given the known roles of hypothalamic ERα in mediating many aspects of physiology, as well as the effect of hypothalamic ERα knockout on cellular tamoxifen response, we sought to ask if hypothalamic loss of ERα would also affect mouse physiological responses to tamoxifen treatment. We therefore measured body temperature, physical activity and bone density in *Esr1*^cKO^ mice treated equivalently and concurrently with wild-type animals (Figure 1A). Interestingly, tamoxifen injection was not associated with any difference in core body temperature in *Esr1*^cKO^ mice compared to vehicle-treated controls (Figure 6A-B). Specifically, we observed no significant differences in heat dissipation or production when comparing tamoxifen to vehicle treatment in *Esr1*^cKO^ mice, as indicated by HLI (Figure 6C-D) and BAT temperature (Figure 6E-F). In contrast to wild-type mice, there was no significant difference in physical activity between tamoxifen and vehicle treatment in *Esr1*^cKO^ mice (Figure 6G-H).

**Figure 6.**
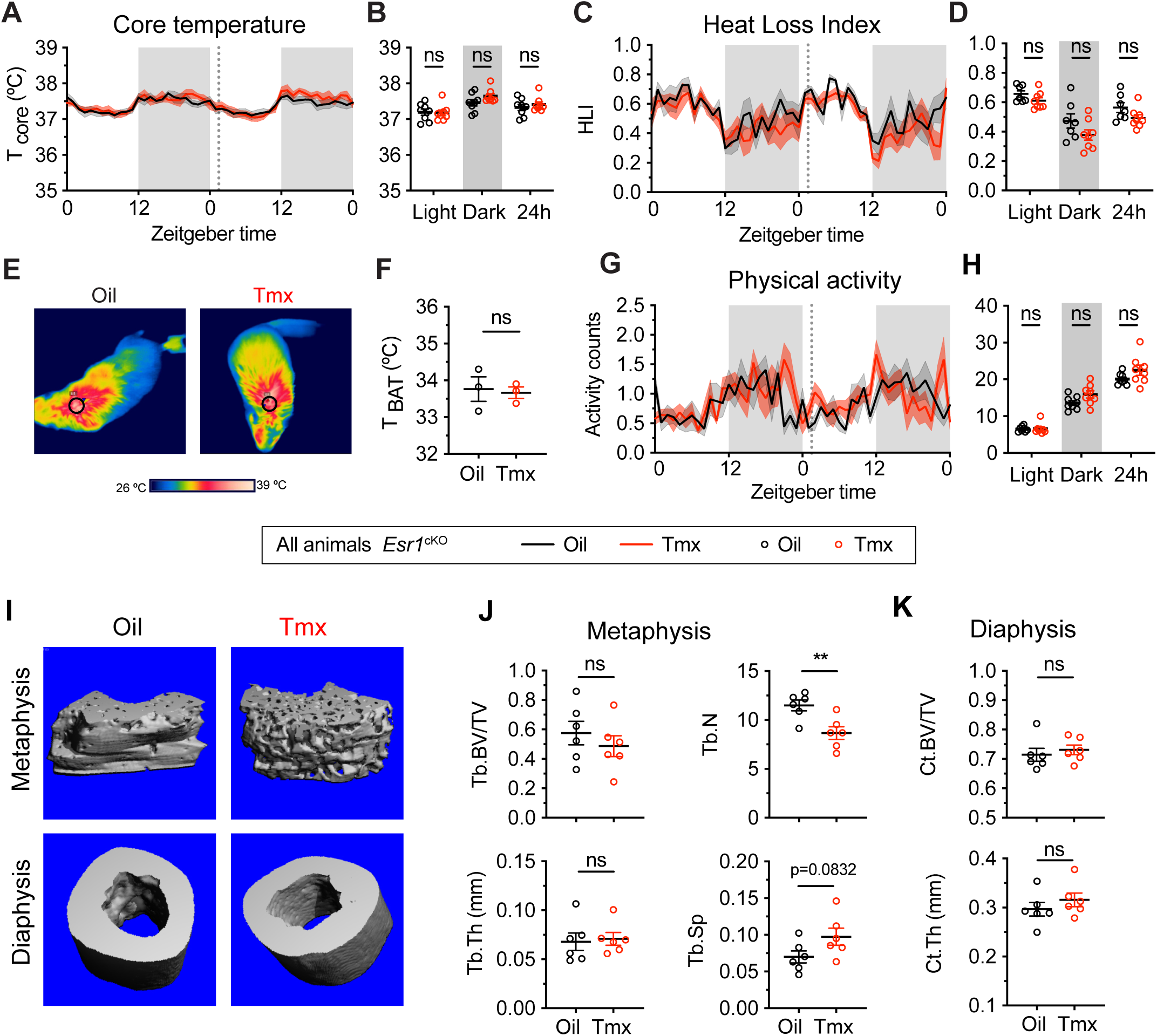
The homeostatic effects of tamoxifen are dependent on hypothalamic ERα. (**A**) Hourly average of core body temperature over 2 days before and 3 weeks after injections (dotted line). (**B**) Average core body temperature from mice shown in panel **A** highlighting per animal averages in light (7:00 to 19:00), dark (19:00 to 7:00), and total 24 h periods (n=8, treatment P=0.7144). (**C**) Heat loss index (HLI) calculated from continuous measurements of core and tail temperature. (**D**) Average HLI from mice shown in panel **C** (n=7-8, P=0.1013). (**E**) Thermographic images of interscapular skin above brown adipose tissues depots in mice injected with either oil or tamoxifen. (**F**) Quantification of temperature of skin above intrascapular BAT depots (n=3, P=0.8048). (**G**) Hourly average of physical activity over 2 days before and after injections (dotted line). (**H**) Average physical activity from mice shown in panel **H** (n=8, P=0.1266). (**I**) Representive microCT images showing bone density of the distal metaphysis and midshaft diaphysis of femurs from tamoxifen or oil treated *Esr1*^cKO^ female mice. (**J**) Trabecular bone volume fraction (BV/TV), trabecular numbersdx (Tb.N), trabecular thickness (Tb.Th) and trabecular separation (Tb.Sp) in distal metaphysis of femurs (n=6). (**K**) Bone volume fraction (Ct.BV/TV) and cortical thickness (Ct.Th) in diaphysis of femurs (n=6). Line shading and error bars represent SEM. ns, non-significant; **, P<0.01 for Sidak’s multiple comparisons test (**B, D** and **H**) or two tailed t-test (**F, J** and **K**).

Knockout of ERα in the hypothalamus largely blocked the effect of tamoxifen on bone physiology as well. No difference was observed in bone volume fraction, thickness or separation of trabeculae when comparing tamoxifen to vehicle treated animals. In contrast to wild-type mice, we observed a tamoxifen-induced reduction in trabecular number (Figure 6I-J). There was no significant difference in bone volume ratio or cortical thickness in the cortical bones of mice treated with tamoxifen compared to mice treated with vehicle (Figure 6I and K). Taken together, these results suggest that the dysregulation of core temperature, reduced movement, and changed bone physiology associated with tamoxifen treatment in wild-type mice are largely dependent upon the expression of ERα in the *Nkx2-1* lineage.

## Discussion

Tamoxifen is a selective estrogen receptor modulator that exerts antiproliferative effects on cancer cells through ERα *(Davies et al. 2011)*, but can exert agonistic or antagonistic effects on the different estrogen receptors depending on cell type and context *(Watanabe et al. 1997, Mo et al. 2013)*. The medial basal hypothalamus is rich in estrogen receptor expression, making it a prime candidate as a tamoxifen target *(Merchenthaler et al. 2004, Gofflot et al. 2007, Matsuda et al. 2013, Saito et al. 2016)*. Here we take the first steps to define the cell types and cellular mechanisms that mediate some of the undesirable side effects experienced by people undergoing tamoxifen therapy. We demonstrate the utility of mice as a good model to study many of tamoxifen’s physiological effects; tamoxifen administration in mice increases heat dissipation, decreases movement, and increases bone density. These effects are similar to those experienced by people receiving tamoxifen therapy, who experience hot flashes, lethargy, and changes in bone density, amongst other symptoms *(Love et al. 1991, Powles et al. 1996, Fisher et al. 1998, Loprinzi et al. 2000, Haghighat et al. 2003, Howell et al. 2005, Kligman et al. 2010, Francis et al. 2015)*.

Using single cell RNA sequencing, we find that tamoxifen alters transcriptomes in the hypothalamus, consistent with the hypothesis that tamoxifen therapy alters central nervous system function *(Eberling et al. 2004, Gibbs et al. 2014, Denk et al. 2015)*. Indeed, tamoxifen and its metabolites are detectable and up to 46-fold higher in the brain than in serum of breast cancer patients *(Lien et al. 1991)*. We report that tamoxifen treatment affects gene expression in many cell types of the hypothalamus, but most strongly in neurons and ependymal cells. These findings are consistent with previous reports of tamoxifen-induced gene expression changes within neurons *(Lopez et al. 2006, Aquino et al. 2016, Sa et al. 2016)*, and tamoxifen-induced stress in neurons and ependymal cells *(Denk et al. 2015)*. Importantly, we detect minor changes in the proportion of different cell types after 28-days of treatment with a clinically relevant dose (0.1 mg/kg) of tamoxifen. This result is in contrast to evidence that the much higher doses used to activate tamoxifen-inducible mouse models (25-100 mg/kg) can inhibit neural progenitor cell proliferation during cortical patterning or induce adipogenesis and prolonged genetic effects *(Ye et al. 2015, Lee et al. 2020)*.

Notably, conditional knockout of the *Esr1* gene encoding ERα did not render the brain insensitive to tamoxifen. Instead it ablated or reversed the direction of a significant number of the tamoxifen induced gene expression changes that were observed in the wild-type hypothalamus. Importantly, this reversal encompassed many functional pathways including those controlling metabolism and neuropeptide signaling in neurons. The many transcriptional changes induced by tamoxifen in *Esr1*^cKO^ mice might be attributed to tamoxifen’s effects on the other estrogen receptors or effects mediated by cells outside of the *Nkx2-1* lineage. Indeed, studies of breast cancer resistance suggest many other signaling pathways, e.g., progesterone receptor, androgen receptor and GPER, may be involved in the actions of tamoxifen on transcription *(Mo et al. 2013, Rondón-Lagos et al. 2016)*. Indeed, progesterone receptor expression in the hypothalamus is regulated by both endogenous estrogens and tamoxifen *(Sa et al. 2016)* and we find that *Pgr* is most abundant in neurons. Although more enriched in mural cells, GPER has been shown to increase acetylcholine release in neurons of the hippocampus *(Gibbs et al. 2014)*. Nevertheless, conditional knockout of ERα in the hypothalamus blocked the majority of the physiological changes observed in wild-type mice by tamoxifen, indicating a pivotal role of ERα in mediating the homeostatic effects of tamoxifen. Together, these data indicate an indispensable role of hypothalamic ERα in the molecular and physiological response to tamoxifen therapy in mice. In addition, our data indicate that tamoxifen exerts potent effects on gene expression in a variety of neuronal subtypes, as well as in ependymal cells. It is very likely that the pleiotropic physiological effects of tamoxifen involve various estrogen responsive cells and hypothalamic regions. Determining which hypothalamic nuclei respond directly to tamoxifen and which nuclei mediate the various effects of tamoxifen will require careful region and cell-type specific *Esr1* deletion studies.

Hot flashes are one of the most common complaints by people undergoing tamoxifen therapy or transitioning to menopause *(Stearns et al. 2002, Kligman and Younus 2010)*. Hot flashes are characterized by frequent, sudden increases of heat dissipation from the skin, often accompanied with sweats and transient decreases in core body temperature *(Stearns et al. 2002)*. The precise mechanisms behind this change in thermoregulation are unknown, however several studies have suggested that a disruption in sex hormone related central control of thermal homeostasis is involved. It has been proposed that this dysregulated heat dissipation is due to a lowered body temperature setpoint, which is influenced by changes in ovarian hormones including estrogens and luteinizing hormone *(Dacks et al. 2010, Dacks et al. 2011, Rance et al. 2013)*. The hypothalamus, a central site for both sex hormone action and body temperature regulation, appears to play a key role in the etiology of hot flashes. Specifically, activating KNDy (kisspeptin, neurokinin B and dynorphin expressing) neurons, which also express ERα, in the ARC increases heat loss from the tail in mice *(Padilla et al. 2018)*. Conversely, ablation of KNDy neurons *(Mittelman-Smith et al. 2012)* or pharmacologically blocking the Neurokinin 3 receptor (NK3R) in the preoptic area (POA) *(Prague et al. 2017, Padilla et al. 2018)* was able to relieve hot flash symptoms in rodents and humans. As tamoxifen readily penetrates the blood brain barrier *(Lien et al. 1991)* and regulates estrogen-responsive neurons in the hypothalamus *(Aquino et al. 2016, Sa et al. 2016)*, it is plausible that tamoxifen modulates these circuits in the hypothalamus to induce or increase the likelihood of hot flashes. Indeed, we detected the highest *Esr1* expression levels in neurons marked by expression of the neurokinin B precursor gene, *Tac2*. Furthermore, *Esr1* expressing neurons in both the ARC and the MPA develop from the *Nkx2-1* lineage and therefore show loss of ERα in *Esr1*^cKO^ mice (Figure 5A). These data are all consistent with the hypothesis that tamoxifen induces hot flashes via estrogen-sensitive neural circuits involved in thermoregulation.

Animal studies indicate that estrogen signaling promotes movement *(Ogawa et al. 2003, Lightfoot 2008)*, although in humans, estrogen related activity changes remain controversial (see review*(Bowen et al. 2011)*). In female breast cancer patients, there is a significant association between ongoing tamoxifen usage and symptoms of fatigue *(Haghighat et al. 2003)*. Our results demonstrate that ERα in the hypothalamus mediates a suppressive effect of tamoxifen on movement. This is consistent with the results that ERα signaling in the hypothalamus regulates physical activity in mice *(Ogawa et al. 2003, Musatov et al. 2007, Xu et al. 2011, Sano et al. 2013, Correa et al. 2015, van Veen et al. 2020)*. Specifically, activation of ERα neurons in the *Nkx2-1* lineage promotes movement *(Correa et al. 2015)* and loss of ERα in the VMH reduces physical activity in mice *(Musatov et al. 2007, Correa et al. 2015)*. In addition, the POA is rich in ERα expression and has been shown to be responsible for estrogen induced running wheel activity *(King 1979, Fahrbach et al. 1985, Takeo et al. 1995)*. Therefore, tamoxifen may act as an ERα antagonist in those estrogen-sensitive hypothalamic regions to suppress physical activity in mice.

Similar to endogenous estrogens, tamoxifen has profound impacts on bone remodeling. Human studies have demonstrated that tamoxifen is protective against bone mineral density loss in after menopause, but induces bone loss before menopause *(Love et al. 1992, Kristensen et al. 1994, Powles et al. 1996, Vehmanen et al. 2006)*. Contrasting this, animal studies have generally shown protective effects of tamoxifen on bone regardless of ovarian function *(Turner et al. 1988, Perry et al. 2005, Starnes et al. 2007)*. Whether these discrepancies are due to fundamental differences between rodent and human bone biology remains to be determined. We found that tamoxifen greatly increased bone mass in wild-type mice, displaying an estrogen-like protective effect. Although this study did not evaluate bone formation and resorption respectively, a number of animal studies have indicated that tamoxifen can stimulate bone formation *(Perry et al. 2005)* and also suppress bone resorption *(Turner et al. 1988, Quaedackers et al. 2001)*, resulting in a net bone accruement. Although different estrogen receptor subtypes may exert distinct effects on bone regulation *(Windahl et al. 2002)*, the action of tamoxifen on bone is likely mediated by ERα but not ERβ *(Quaedackers et al. 2001)*. Within the hypothalamus, depletion of ERα in the medial basal hypothalamus or specifically within the KNDy neurons results in impressive increases in bone mass *(Farman et al. 2016, Herber et al. 2019)*. Thus it is possible that any effects of tamoxifen on bone that are masked by the already altered bone metabolism in *Esr1*^cKO^ mice. Nevertheless, we observed no effect of tamoxifen on bone volume in *Esr1*^cKO^ mice, implying that tamoxifen’s effects on bone are at least partially mediated by ERα expression within the *Nkx2-1* lineage.

Although the *Nkx2-1* lineage includes cells in the thyroid, pituitary and lung, *Esr1* expression is not altered in these peripheral tissues of *Esr1*^cKO^ mice *(Herber et al. 2019)*, leaving the hypothalamus as the most likely mediator of the ERα-dependent physiological effects of tamoxifen reported here. Despite this, it is very likely that peripheral tissues are also affected by systemic tamoxifen administration, leading to other physiological effects that we did not examine. Indeed, there is evidence that estrogen suppression therapies, including tamoxifen or aromatase inhibitors, can lead to the dysregulation of energy balance and glucose homeostasis in humans and when modeled in rodents *(Lopez et al. 2006, Lampert et al. 2013, Ceasrine et al. 2019, Scalzo et al. 2020)*. Together, these studies reveal widespread effects of tamoxifen therapy on the hypothalamus and implicate ERα signaling within the *Nkx2-1* lineage as a major mediator of the effects of tamoxifen on thermoregulation, physical activity, and bone homeostasis.

In summary, the rodent model of tamoxifen therapy provided here mimics several of the key side effects of endocrine therapy observed in humans and highlights a central mechanism for understanding these specific adverse outcomes. A recent survey conducted in the online breast cancer communities showed that women often experienced more of tamoxifen’s side effects than men and one-third of patients did not feel that their side effects were taken seriously *(Berkowitz et al. 2020)*. Our study highlights many ways in which tamoxifen treatment could impact quality of life in breast cancer patients and takes the first step towards understanding the mechanisms of these side effects in order to develop better treatments.

## Acknowledgements

This research was supported by V Scholar Awards (V2017-007 and DVP2020-005) from the V Foundation for Cancer Research to SMC, NIH R21CA249338 to SMC, Pilot Awards from the Iris Cantor-UCLA Women’s Health Center/UCLA National Center of Excellence in Women’s Health and NIH National Center for Advancing Translational Science (NCATS) UCLA CTSI (UL1TR001881) to ZZ, JEV, and SMC. ZZ was supported by an American Heart Association Postdoctoral Fellowship (18POST33960457) and GD was supported by NIEHS NIH T32ES015457.

## Methods

### Mice

All mice were maintained under a 12:12 h L/D schedule at room temperature (22 °C ∼ 23 °C) and provided with food and water *ad libitum*. Mice expressing the NK2 homeobox transcription factor 1 (*Nkx2-1*; also known as *Ttf1*) Cre driver transgene (Tg(*Nkx2-1*-*Cre*)2Sand) and the *Esr1* floxed allele (*Esr1*tm1Sakh) were maintained on a CD-1;129P2 mixed background. We cross-bred mice carrying *Nkx2-1 Cre* with mice expressing *Esr1 flox* to generate ERα specific knock-outs in the hypothalamus. *Cre*-negative littermates were selected as controls. A total of 50 female mice were used. Mice were 8-10 weeks old at the start of the experiments. All studies were carried out in accordance with the recommendations in the Guide for the Care and Use of Laboratory Animals of the National Institutes of Health. UCLA is AALAS accredited and the UCLA Institutional Animal Care and Use Committee (IACUC) approved all animal procedures.

### Tamoxifen administration

Tamoxifen (Sigma) was dissolved first in ethanol and then diluted in corn oil at a final concentration of 100 μg/mL and 0.5% of ethanol. Accordingly, the vehicle was prepared in corn oil containing 5% ethanol. We performed daily subcutaneous injection of tamoxifen at a dose of 0.1 mg/kg or an equal volume of vehicle control for 4 weeks. We estimated that this dose models human tamoxifen exposure based on studies of tamoxifen in human serum, estimates of total blood volume in mice, and bioavailability of tamoxifen delivered via subcutaneous injection*(Slee et al. 1988)*. Additionally, it is on the low end of what is used effectively in previous studies*(Wade and Heller 1993, Perry et al. 2005)*, which is desirable to minimize off-target effects.

### Telemetry recording

Mice were anaesthetized with isofluorane and received combinatorial analgesics (0.01 mg/mL buprenorphine, 0.58 mg/mL carprofen) pre- and post-any surgeries. A G2 eMitter (Starr Life Sciences) was implanted to the abdominal cavity and attached to the inside body wall of a mouse. Mice were single-housed in cages placed on top of ER4000 Energizer/Receivers. Gross movement and core body temperature were measured every 5 min using VitalView software (Starr Life Sciences). These measurements were collected continuously for 3 days on and 3 days off for the duration of the 28-day period. Telemetry data show 24-hour averages from the 2 telemetry sessions before and 3 weeks after tamoxifen or vehicle injections. Tail skin temperature was monitored every 5 min using a Nano-T temperature logger (Star-Oddi) that was attached to the ventral surface and 1 cm from the base of the tail in a 3D-printed polylactic acid collar modified from Krajewski-Hall et al*(Krajewski-Hall et al. 2018)*. Tail skin data show 24-hour averages for the same time periods as the telemetry data.

### Thermal imaging

Infrared thermal images were captured using e60bx thermogenic camera (FLIR Systems) and analyzed using the FLIR Tools software. All images were obtained at a constant distance to subject in awake animals at light phase. BAT skin temperature was defined from the average temperature of a spherical area centered on the interscapular region.

### Single-cell dissociation and library preparation

To avoid circadian differences and batch effects, all four groups were performed in parallel at the same time across 4 days with n=4 mice per day. Mice were euthanized two hours after the last administration of tamoxifen or vehicle. The brain was freshly collected into ice-cold HABG buffer (Fisher Scientific, Hampton, NH, USA) containing freshly prepared papain. A coronal section containing the hypothalamus was quickly dissected along the boundary of rostral and caudal spherical groves from the bottom of the brain. A third cut was made along the white fiber of anterior commissure to get a square block of hypothalamic tissue (Figure 3 – Supplement 1A). The hypothalamus tissue was then disassociated and prepared into a single-cell suspension at a concentration of 100 cells/μl in 0.01% BSA-PBS using a previously described protocol *(Liu et al. 2020)*. Briefly, dissected tissues were incubated in a papain solution for 30 min at 30 °C then washed with HABG. Using a siliconized 9-in Pasteur pipette with a fire-polished tip, the cells were triturated carefully to help dissociate the tissue. Next, to separate the cells and remove debris, the cell suspension was placed on top of a prepared OptiPrep density gradient (Sigma Aldrich, St. Louis, MO, USA) then centrifugated at 800 g for 15 min at 22 °C. After debris removal, the cell suspension containing the desired cell fractions was centrifugated for 3 min at 22 °C at 200 g, and the supernatant was discarded. Finally, the cell pellet was re-suspended in 0.01% BSA (in PBS) and filtered through a 40-micron strainer (Fisher Scientific, Hampton, NH, USA). The cells were then counted and diluted to appropriate cell density.

Barcoded single cells, or STAMPs (single-cell transcriptomes attached to microparticles), and cDNA libraries were prepared as described previously *(Liu et al. 2020)* and in line with the online protocol v3.1 from McCarroll Lab (http://mccarrolllab.org/download/905/). Briefly, to generate STAMPS, the prepared single cell suspensions, EvaGreen droplet generation oil (BIO-RAD, Hercules, CA, USA), and ChemGenes barcoded microparticles (ChemGenes, Wilmington, MA, USA) containing unique molecular identifiers (UMIs) and cell barcodes were co-flowed through a FlowJEM aquapel-treated Drop-seq microfluidic device (FlowJEM, Toronto, Canada) at recommended flow speeds (oil: 15,000 μl/h, cells: 4000 μl/h, and beads 4000 μl/h). After breakage of the droplets, the beads were washed and suspended in reverse transcriptase solution. Prior to PCR amplification, samples underwent an exonuclease I treatment. Next, the beads were washed, counted and aliquoted into PCR tubes (6000 beads/tube) for PCR amplification (4+11 cycles). The cDNA quality was checked using a High Sensitivity chip on BioAnalyzer (Agilent, Santa Clara, CA, USA) then fragmented using Nextera DNA Library Preparation kit (Illumina, San Diego, CA, USA) with multiplex indices. The libraries were then purified, quantified, and sequenced on an Illumina HiSeq 4000 (Illumina, San Diego, CA, USA).

### Analysis of single-cell RNA-seq

Single cell transcriptomic data were analyzed in R version 3.6.1 using the package “Seurat” version 3.2.0 and custom Seurat helper package “ratplots” version 0.1.0 written for this study. All custom functions and complete analysis scripts are available at https://github.com/jevanveen/. Cells were filtered for quality with the following criteria: cells with > 15% of reads coming from mitochondrial genes were excluded and cells with fewer than 200 or more than 4000 genes detected were excluded. Samples derived from the four experimental conditions: Wild-type, vehicle treated; Wild-type, tamoxifen treated; *Esr1*^*cKO*^, vehicle treated; and *Esr1*^*cKO*^, tamoxifen treated, were aligned based on the expression of 2000 conserved highly variable markers in each group. Cells were clustered based on transcriptome similarity, using a shared nearest neighbor algorithm *(Waltman et al. 2013)*. For each cell cluster, marker genes were determined by comparing expression in the given cluster against all other clusters using the smart local moving algorithm to iteratively group clusters together *(Blondel et al. 2008)*. For all differentially expressed gene marker analyses, statistical significance testing was performed with the Seurat default Wilcoxon rank-sum-based test and Benjamini–Hochberg method for multiple-testing correction. Gene set enrichment analyses were performed in R using the package “fgsea” version 1.14.0. GSEA analyses to find estrogen signatures were done using the gene ontology pathway “GO_ESTROGEN_RESPONSE” taken from mSigDB version 6.2. For general pathway analyses, a custom set of pathways was assembled from mSigDB “Hallmark” pathways version 6.2 and selected GO pathways (version 6.2) particularly relevant to neural development and function. The complete list of these pathways is given in Table S2.

Cells annotated as neurons were extracted and re-clustered using the same steps used to cluster all cells, with the following differences: 500 highly variable genes were used to align neurons from different treatment groups. No additional quality filtering was performed. Neuronal sub-types were annotated based on top marker genes, defined as the highest logFC compared to all other neuronal sub-types, whilst meeting the criterion of adjusted P value < .05. Gene set enrichment analyses were performed on neuronal sub-types using the same gene sets and methods performed for general cell types. All graphs and tables in figures 3-5 were produced using the R packages “ggplot2” version 3.3.2 and custom functions in “ratplots” version 0.1.0.

### Micro-computed tomography

Left femoral bones from the hind legs were dissected, cleaned of any soft tissue and frozen in PBS at −20 °C till further analysis. Samples were scanned in 70% ethanol using Scanco Medical μCT 50 specimen scanner with a voxel size of 10mm, an X-ray tube potential of 55 kVp and X-ray intensity of 109 μA. Scanned regions included 2 mm region of the femur proximal to epiphyseal plate and 1 mm region of the femoral mid-diaphysis. For the analysis, a trabecular bone compartment of 1mm length proximal to the epiphyseal plate was measured. Cortical parameters were assessed at the diaphysis in an adjacent 0.4 mm region of the femur. In both specimen and in vivo scanning, volumes of interest were evaluated using Scanco evaluation software. Representative 3D images created using Scanco Medical mCT Ray v4.0 software.

### Immunohistochemistry

Mice were perfused transcardially with ice-cold PBS (PH=7.4) followed by 4% paraformaldehyde (PFA) in PBS. Brains were post fixed in 4% PFA overnight, dehydration in 30% sucrose for 24 hours, embedded in OCT and stored in −80 °C before sectioning. Coronal sections were cut under cryostat (Vibratome) into 8 equal series at 18 μm. ERα immunoreactivity was detected using hot based antigen retrieval immunohistochemistry protocol. Briefly, sections were first incubated in antigen retrieval buffer (25 mM Tris–HCl, 1 mM EDTA, and 0.05% SDS, pH 8.5) at 95 °C for 40 min. Sections were then blocked for 1 h in 10% BSA and 2% normal goat serum (NGS) and incubated overnight at 4 °C with primary antibody (ERα, 1:250, sc-8002, Santa Cruz). Following 3 × 10 min washing in PBS, sections were incubated with Fluorophore conjugated goat anti-mouse secondary antibody (Thermo Fisher Scientific) for 2 h at room temperature. After washing, section were incubated with DAPI, PBS washed and coverslipped with Fluoromount-G.

The images were taken by DM1000 LED fluorescent microscope (Leica). Cyan/magenta/yellow pseudo-colors were applied to all fluorescent images for color-friendly accessibility. Image processing was performed using the Leica Application Suite (Leica) and ImageJ (NIH).

### RNA isolation and Real Time PCR (qPCR)

Interscapular BAT was snap frozen in liquid nitrogen and stored at −80 °C until analysis. Total RNA from BAT was isolated using Zymo RNA isolation kit (ZYMO Research) and RNA yield was determined using a NanoDrop D1000 (Thermo Fisher Scientific). cDNA synthesis was performed with equal RNA input using the Transcriptor First Strand cDNA synthesis kit (Roche Molecular Biochemicals). Quantitative PCR was performed using C1000 Touch Thermal Cycler (BioRed) and SYBR mix (Bioline, GmbH, Germany), using validated primer sets *(Herber et al. 2019)* (Table 1).

## Statistics

Data are represented as mean ± standard error of the mean (SEM). Data with normal distribution and similar variance were analyzed for statistical significance using two-tailed, unpaired Student’s t-tests. Time course data were analyzed by repeated measures two-way ANOVA or mixed model followed by Sidak’s multiple comparisons. Significance was defined at a level of *P* < 0.05. Statistics were performed using GraphPad Prism 8 and RStudio.

---

**Figure 1 - Supplement 1.**
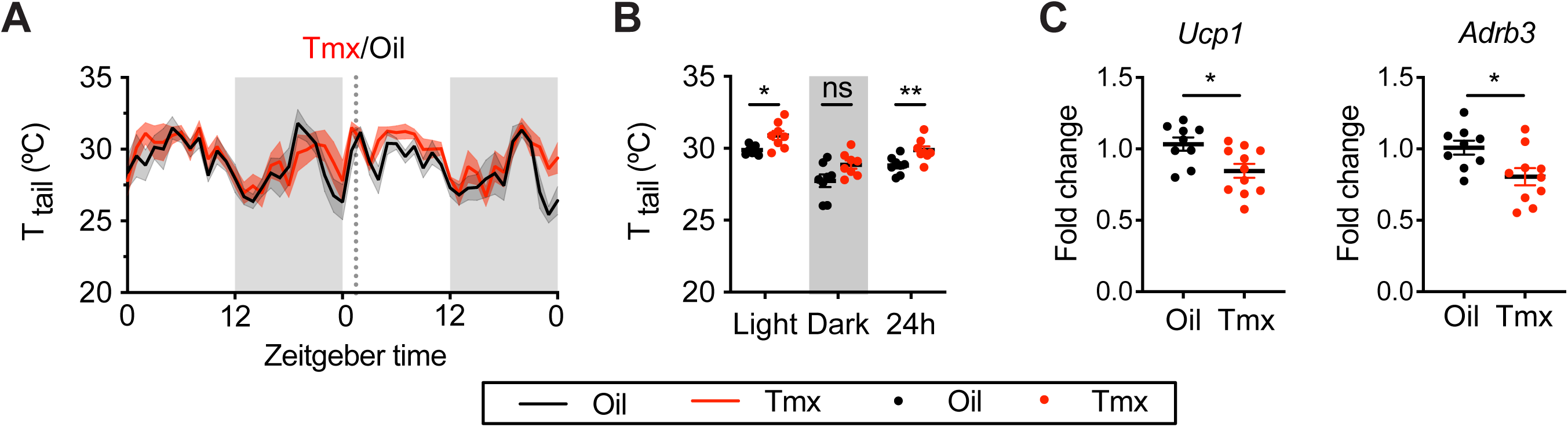
Tamoxifen increased temperature in tail and suppressed thermogenic gene expression in BAT. (**A**) Hourly average of tail skin temperature over 3 days before and 3 weeks after injections (dotted line) measured every 5 min using attached thermal logger. (**B**) Average tail temperature from mice shown in panel **A** highlighting per animal averages in light, dark, and total 24 h periods (n=8, treatment **). (**C**) Relative mRNA expression of *Ucp1* and *Adrb3* from BAT extracts (n=8-10). ns, non-significant; *, P<0.05; **, P<0.01 for Sidak’s multiple comparison tests (**B**) following a significant effect of treatment in a two-way RM ANOVA or two tailed t-tests (**C**).

**Figure 3 - Supplement 1.**
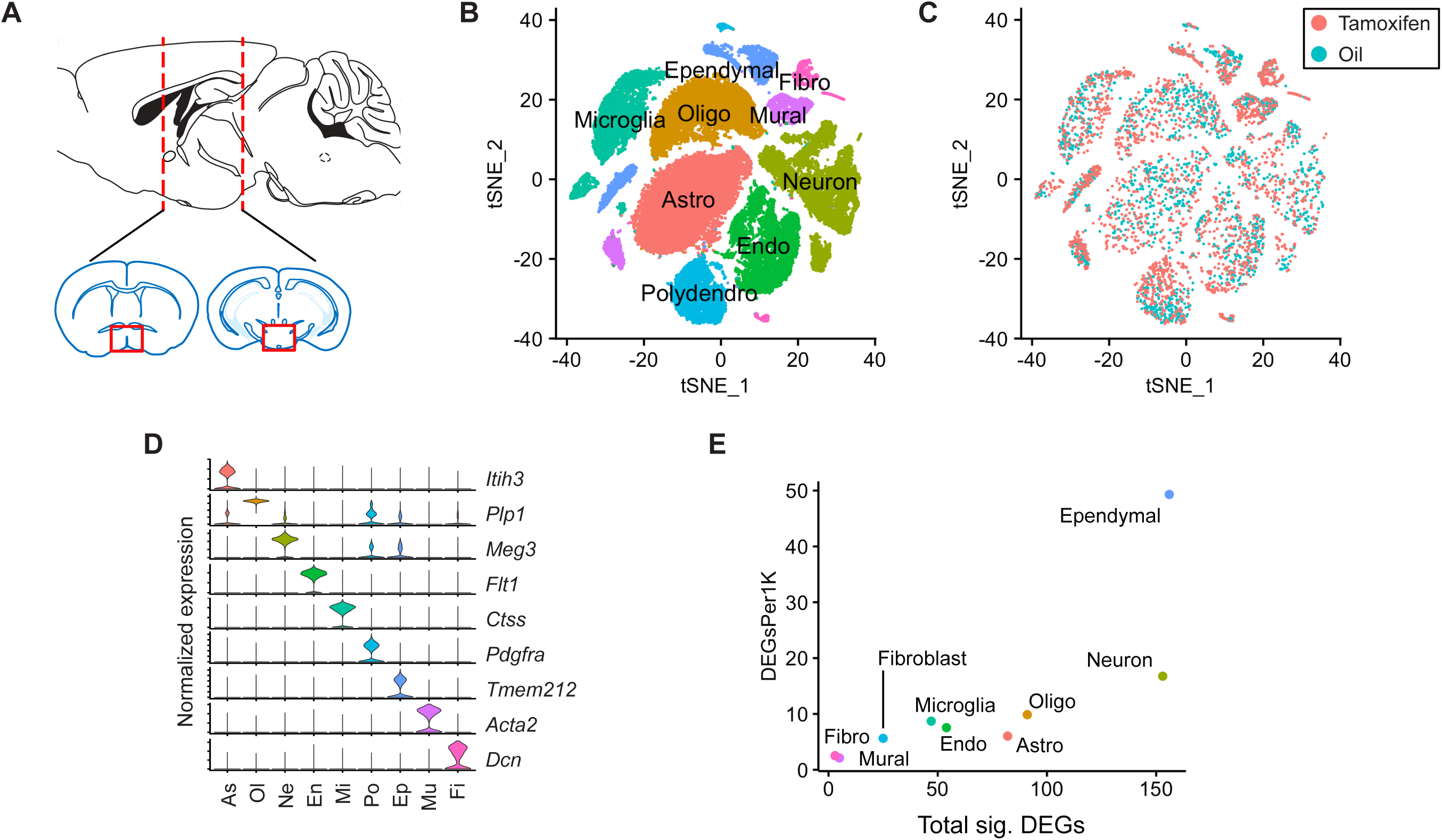
Tamoxifen treatment induces gene expression changes in various cell types of the hypothalamus. **(A)** Schematic of the hypothalamic area dissected for scRNA-seq (Bregma +0.4 to –3 mm). (**B**) tSNE showing clustering of the major cell types of the hypothalamus based on single cell transcriptomics. (**C**) tSNE of hypothalamic cells, comparing cells derived from female mice receiving 28 daily tamoxifen injections (pink) or 28 daily oil injections (blue). (**D**) Violin plots showing normalized expression of cluster defining markers in all clusters. (**E**) Plot comparing the total number of significant (Bonferroni adj. P < .05) tamoxifen regulated genes to the number of significant tamoxifen regulated genes per thousand cells in each cluster. All analyses done from cells harvested from female mice injected daily with oil (n=3) or tamoxifen (n=5) over 28 days.

**Figure 4 - Supplement 1.**
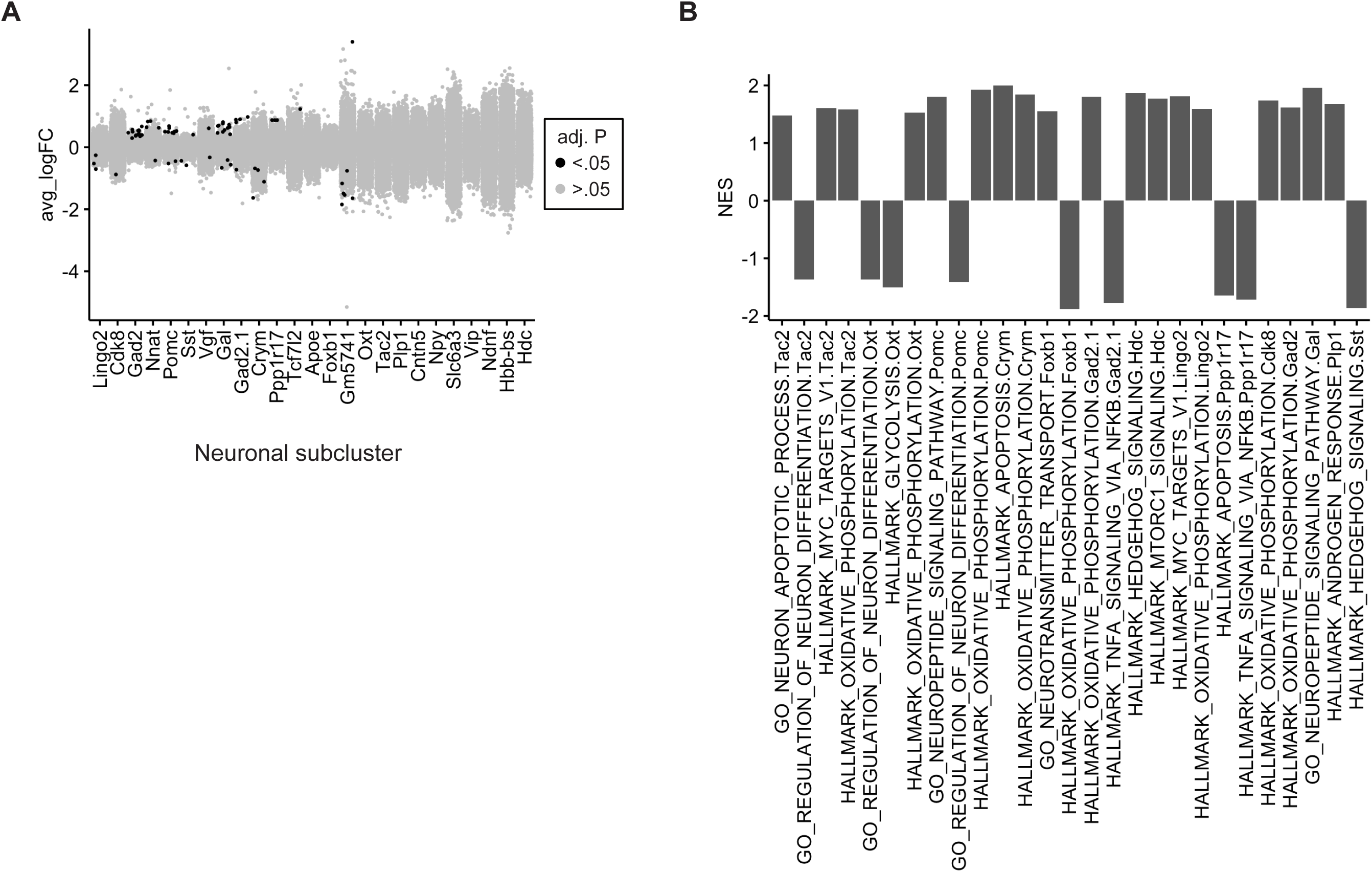
Tamoxifen induced gene expression changes in neurons of the hypothalamus. (**A**) Collapsed volcano plot of differences in gene expression associated with tamoxifen treatment in the various neuronal sub-types. (**B**) GSEA pathways significantly (adj. P < .05) upregulated or downregulated by tamoxifen in individul neuronal subclusters

**Figure 5 - Supplement 1.**
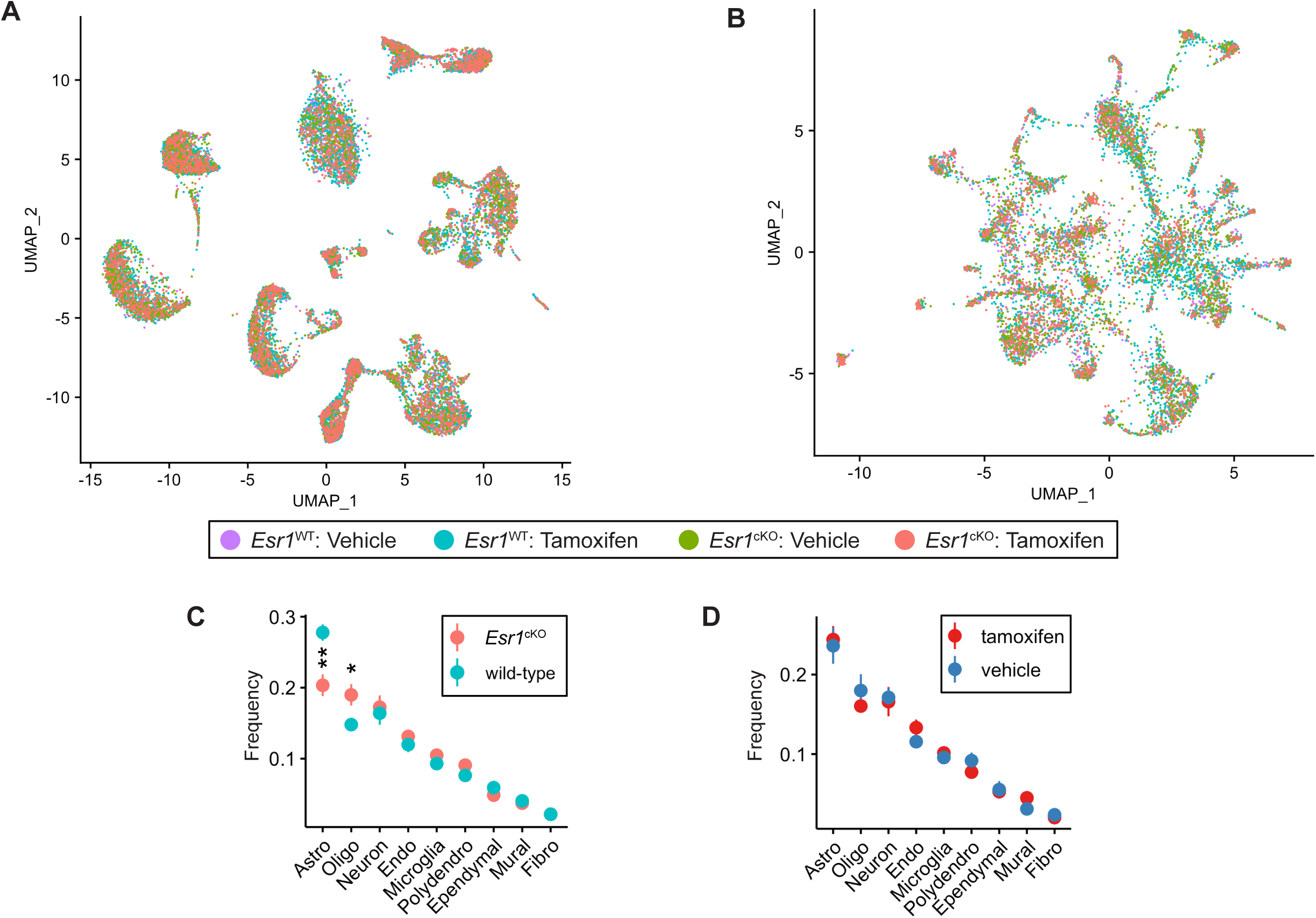
identification of distinct cell clusters in 4 aligned treatment groups. (**A**) UMAP of all hypothalamic cell types showing the four studied treatment groups. (**B**) UMAP of hypothalamic neuronal types showing the four studied treatment groups. (**C**,**D**) Proportion of each cell type recovered, comparing *Esr1*^cKO^ to wild-type **C** and tamoxifen to vehicle **D**. Two way anova shows effect of *Esr1* genotype on proportion of astrocytes (F = 13.3 adj. P = .003) and oligodendrocytes (F = 7.8, adj. P = .016), but no effect of tamoxifen treatment on proportion of any cell type.

**Figure 5 - Supplement 2.**
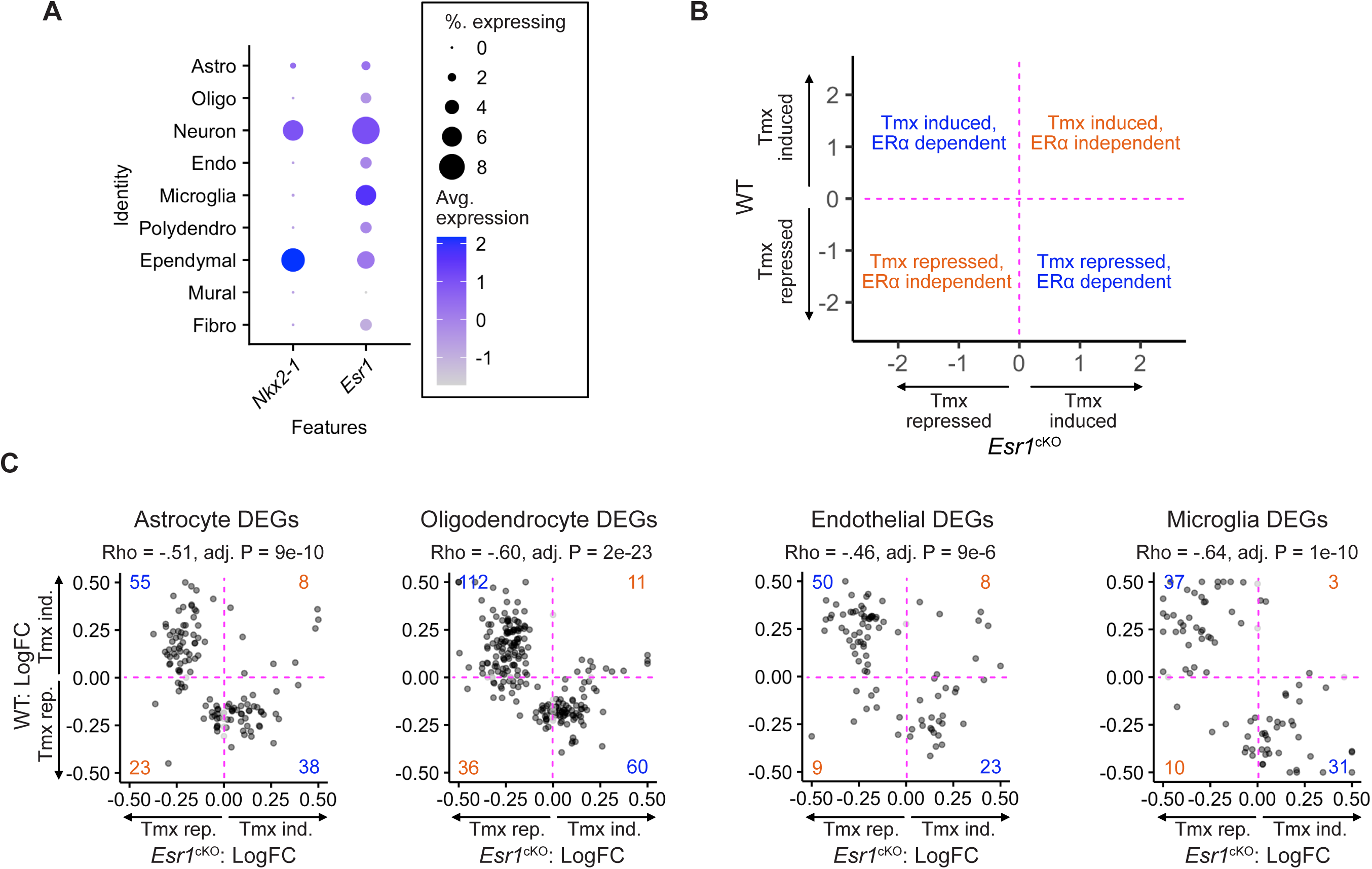
Comparison of gene expression changes by tamoxifen in WT and *Esr1*^cKO^ mice. (**A**) Dotplot showing expression of *Esr1* and *Nkx2-1* in the major cell types of the hypothalamus. (**B**) Key to comparison of genes induced by tamoxifen in wild-type cells compared to how they are regulated by tamoxifen in *Esr1*^cKO^ animals. (**C**) Gene by gene comparisons of how tamoxifen treatment affects expression in wild-type and *Esr1*^cKO^ cells. Only genes significantly up or downregulated by tamoxifen in either wild-type or mutant cells are shown.

**Figure 5 - Supplement 3.**
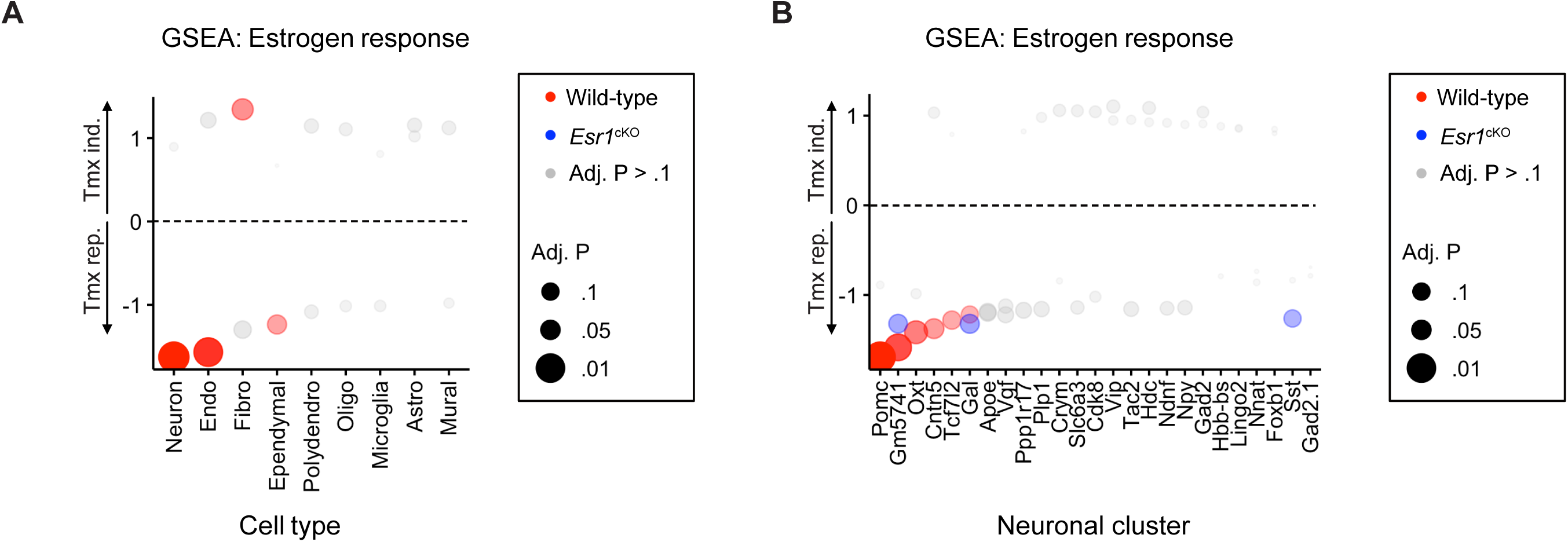
Comparison of gene expression changes by tamoxifen in WT and *Esr1*^cKO^ mice. (**A**) GSEA normalized enrichment scores (NES) showing tamoxifen regulation of estrogen responsive genes in wild-type and *Esr1*^cKO^ hypothalamic cell types. (**B**) GSEA normalized enrichment scores (NES) showing tamoxifen regulation of estrogen responsive genes in wild-type and *Esr1*^cKO^ neuronal sub-types.

